# TSUMUGI: a platform for phenotype-driven gene network identification from comprehensive knockout mouse phenotyping data

**DOI:** 10.64898/2026.02.18.706720

**Authors:** Akihiro Kuno, Kinari Matsumoto, Taito Taki, Satoru Takahashi, Seiya Mizuno

## Abstract

**Summary:** Deciphering complex organismal phenotypes requires elucidation of coordinated functions among multiple genes, yet this remains a fundamental challenge in functional genetics. The International Mouse Phenotyping Consortium (IMPC) has recently established a comprehensive phenotypic atlas based on systematic single-gene knockout mouse lines, providing an unprecedented resource for gene-phenotype associations in mammals. However, extracting phenotype-associated multiple gene relationships remains challenging. Here, we present TSUMUGI, a platform that identifies a phenotype-driven gene network by leveraging gene-phenotype associations from the IMPC. TSUMUGI enables users to initiate analyses from a user-specified phenotype or gene list and interactively explore a gene network. In addition, TSUMUGI highlights human disease-associated genes, supports flexible network filtering, and allows seamless export of a gene network for downstream analyses. By linking shared phenotypic signatures to putative functional gene modules, TSUMUGI provides a framework for systematic interpretation and hypothesis generation of complex organismal phenotypes.

**Availability and implementation:** The web application is available online at https://larc-tsukuba.github.io/tsumugi/. The command-line tools are distributed via PyPI (https://pypi.org/project/TSUMUGI) and Bioconda (https://bioconda.github.io/recipes/tsumugi/README.html). Source code and documentation are hosted at GitHub (https://github.com/akikuno/TSUMUGI-dev) and archived on Zenodo (https://doi.org/10.5281/zenodo.18464478) under the MIT license.

## 1. Introduction

Complex organismal phenotypes are often driven by coordinated interactions among multiple genes (Costanzo et al., 2019). Although multiple gene associations have been extensively investigated using protein-protein interactions (PPI), biological pathways, and gene co-expression networks, these approaches primarily focus on molecular and intracellular relationships and may not fully reflect gene interactions at the organismal level (Allayee et al., 2023). Similarly, genome-wide association studies and polygenic risk score approaches identify genetic risk factors for complex traits but do not explicitly capture coordinated gene interactions that give rise to organismal phenotypes (Lewis and Vassos., 2020).

The International Mouse Phenotyping Consortium (IMPC) has generated comprehensive phenotypic profiles for more than 9,000 single-gene knockout (KO) mouse lines, providing a large-scale standardized resource for systematic gene-phenotype analyses (Dickinson et al., 2016; Wilson et al., 2025). This dataset enables direct comparison of organism-wide phenotypic consequences resulting from individual gene disruptions. However, because the IMPC dataset is fundamentally organized around single-gene knockout lines, extracting phenotypic relationships across multiple genes remains challenging.

Here, we present TSUMUGI (Trait-driven Surveillance for Mutation-based Gene module Identification), a platform designed to identify a phenotype-driven gene network from the IMPC dataset. TSUMUGI is based on the working hypothesis that genes whose knockout mice exhibit similar phenotypes participate in shared biological systems, and accordingly quantifies phenotypic similarity between KO mouse lines. By converting gene-phenotype associations into a phenotype-driven gene network, TSUMUGI enables systematic identification of gene associations underlying organism-level phenotypes.

TSUMUGI supports flexible analysis workflows initiated from a user-defined phenotype or a gene list. Users can interactively explore a phenotype-driven gene network, apply customizable filtering criteria, highlight human disease-associated genes, and perform centrality analyses. The extracted gene network can be readily exported for downstream analyses and integration with external tools.

TSUMUGI is available as both a web application and a command-line interface. The web application (https://larc-tsukuba.github.io/tsumugi) provides an intuitive environment for exploratory analysis, whereas the command-line interface, distributed via PyPI and Bioconda, supports reproducible, large-scale analyses with advanced filtering options.

## 2. Methods

### 2.1 Data Sources

Gene-phenotype association dataset derived from single-gene knockout mouse lines was obtained from the IMPC statistical summary table (https://ftp.ebi.ac.uk/pub/databases/impc/all-data-releases/release-23.0/results/, statistical-results-ALL.csv.gz, release: 23.0). From this dataset, the MGI marker symbol, Mammalian Phenotype Ontology (MP) term, effect size, P-value, sex-specific P-value, zygosity, pipeline name, and procedure name were extracted for each gene-phenotype record.

Life-stage annotations were assigned to each gene-phenotype record as follows. An “Embryo” label was assigned when the procedure included embryonic screening (E9.5, E10.5, E12.5, Embryo LacZ, E14.5, E14.5-E15.5, or E18.5). An “Interval” (17 weeks to 48 weeks) or “Late” (49 weeks or later) label was assigned when the pipeline name contained “interval” or “late”, respectively; otherwise, an “Early” label was assigned.

Mouse orthologues of human disease-associated genes were obtained from the IMPC Disease Models Portal (https://diseasemodels.research.its.qmul.ac.uk/, impc_phenodigm.csv, release: 2025-05-15) (Cacheiro et al., 2024). In the downloaded table, the description column contains mouse allele symbols, zygosity, and life-stage information. This field was normalized to match the phenotype dataset and appended accordingly.

MP annotation files were downloaded from OBO Foundry (https://obofoundry.org/ontology/mp.html, mp.obo, release: 2025-08-27) (Jackson et al., 2021) and used to calculate phenotype similarity.

### 2.2 Extraction of abnormal phenotypes

In the IMPC statistical summary table, an MP term is assigned to a gene-phenotype record when either the P-value or the sex-specific P-value is below 0.00001. We therefore extracted records with an assigned MP term as abnormal phenotypes for downstream analyses. Sex specificity was defined as “Female,” “Male,” or “None” according to the significance of sex-specific P-values.

### 2.3 Phenotype similarity metrics

Phenotypic similarity between KO mouse lines was calculated based on the Phenodigm algorithm (Smedley et al., 2013).

Term similarity for each pair of MP terms was defined as the geometric mean of the Resnik score and the Jaccard index. Information content (IC) was calculated from the number of descendant terms and used to compute the Resnik score. Terms with IC below the 5th percentile were considered overly general, and their Resnik scores were set to zero. The Jaccard index was calculated as the overlap ratio between the parent term sets of the MP terms.

A weighted similarity matrix was then constructed for all pairwise combinations of MP terms associated with two KO mouse lines by multiplying the term similarity score by a metadata concordance score. The metadata concordance score was defined as 1.0 when genotype, sex, and life stage were fully matched, and as 0.75, 0.5, or 0.25 depending on the number of mismatched metadata attributes.

Finally, Phenodigm-based scaling was applied to the weighted similarity matrix, yielding a phenotype similarity score between 0 and 100 for each gene pair.

### 2.4 Network construction and visualization in the web application

Gene pairs sharing at least three abnormal phenotypes were extracted, with each gene represented as a node in an undirected graph. The network was visualized using Cytoscape.js (version 3.30.4) (Franz et al., 2015).

## 3. Results

### 3.1 Web application

TSUMUGI provides three types of entry points: Phenotype, Gene, and Gene List, through which users can submit a MP term or gene symbols of interest (Fig. 1A). Upon submission, a dashboard comprising a network view with control panels is displayed (Fig. 1B).

**Figure 1.**
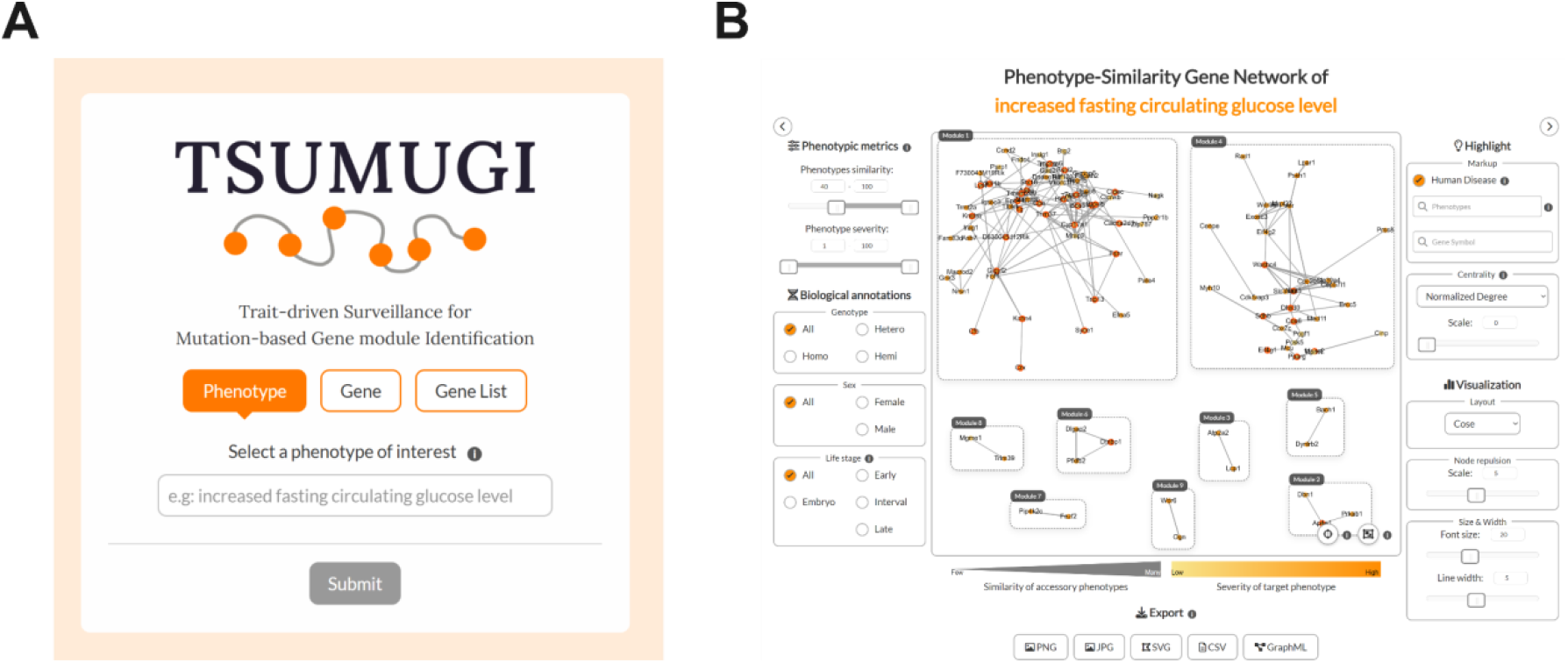
TSUMUGI web interface. (A) Top page. TSUMUGI accepts a phenotype term, gene symbol, or gene list as input. (B) Network page. The central panel displays the phenotype-driven gene network. The left panel provides network filtering options, the right panel provides highlighting and visualization options, and the bottom panel provides export functions.

On the top page, when the Phenotype entry is selected, TSUMUGI identifies genes whose KO mice exhibit the user-specified phenotype and groups them based on phenotype similarity. In contrast, the Gene entry identifies genes whose KO phenotypes resemble those of the user-specified gene. The Gene List entry groups genes within the submitted gene list (Fig. S1).

In the network view, nodes represent genes and edges represent phenotype similarity. Node colour intensity indicates phenotype severity, while edge thickness reflects the phenotype similarity score. The subnetworks are displayed as modules corresponding to connected components, where genes within a module share high phenotypic similarity, while similarity between modules is low. By selecting a module, users can inspect a list of phenotypes enriched within that module. This feature allows users to assess which phenotypes each module contributes to and to iteratively narrow down modules of interest (Fig. S2).

The left and right control panels provide interactive filtering based on similarity scores, severity, and metadata (zygosity, sex, and life stage), as well as highlighting functions based on human disease-association, phenotypes or genes, and network centrality. All interactions are immediately reflected on the client side.

The gene network can be exported as static images (PNG, JPG) and vector graphics (SVG). In addition, module-wise gene lists and associated phenotype information can be exported in CSV and GraphML formats, enabling downstream analyses using external tools.

### 3.2 Command-line interface

In addition to the web application, TSUMUGI provides a command-line interface. This allows researchers to process IMPC phenotype data in a local environment and construct a custom network. Compared with the web tool, the command-line interface enables more flexible extraction of a phenotype similarity network based on phenotypes, genes, and metadata.

Using the command-line tool, users can extract or exclude specific gene pairs based on phenotype information and metadata (Table 1). For example, the ‘tsumugi mp’ command enables extraction or exclusion of gene pairs associated with specified MP terms. This functionality is useful, for instance, when extracting the gene network that is associated with diabetes-related phenotypes but not with reproductive system phenotypes.

**Table 1.** Summary of supported operations available in the TSUMUGI suite.

| Utility | Description |
| --- | --- |
| run | Build the TSUMUGI network locally from the IMPC dataset |
| mp --include/--exclude | Retain or remove gene pairs that include or do not exhibit the specified MP term and its descendant terms |
| zygosity --keep/--drop | Retain or remove gene pairs associated with specified zygosity types (Homo/Hetero/Hemi) |
| sex --keep/--drop | Retain or remove gene pairs associated with specified sex categories (Male/Female/None) |
| life-stage --keep/--drop | Retain or remove gene pairs associated with specified life stages (Embryo/Early/Interval/Late) |
| count --min/--max | Filter gene pairs or genes by the number of associated phenotypes |
| score --min/--max | Filter gene pairs by phenotype similarity score |
| genes --keep/--drop | Retain or remove specified genes |
| build-graphml | Export the network as GraphML format |
| build-webapp | Export the network as TSUMUGI web application assets |

In particular, the ‘tsumugi mp --exclude’ command allows exclusion of gene pairs that are not associated with a specified MP term and its descendant terms. Conventionally, the absence of phenotype annotations for a gene made it unclear whether the phenotype was unmeasured or not significant. By leveraging IMPC-provided information indicating that a phenotype was measured but not significant, TSUMUGI enables exclusion of non-significant phenotypes from the analysis.

Furthermore, as in the web application, the command-line tool supports export in GraphML format, facilitating straightforward integration of TSUMUGI outputs into other analysis pipelines and workflows.

## 4. Discussion

TSUMUGI enables the identification of putative gene interactions associated with organism-level phenotypes. Through flexible filtering options, users can extract gene modules tailored to specific research questions. For example, by restricting the analysis to female- or male-specific phenotypes, users can identify sex-dependent gene modules underlying a phenotype of interest, thereby facilitating the exploration of sexually dimorphic gene interactions. Similarly, by focusing on phenotypes observed at late life stages (49 weeks or later), TSUMUGI allows the identification of gene modules potentially involved in aging-related traits. Together, these features position TSUMUGI as a hypothesis-generating framework for linking shared phenotypic signatures to coordinated gene functions.

A representative study related to TSUMUGI integrated gene and phenotype networks based on pre-defined interactions such as PPI and OMIM to prioritize candidate gene-phenotype associations (Li and Patra., 2010). In contrast, TSUMUGI constructs a gene network directly from large-scale KO mouse phenotype data without relying on pre-defined interaction maps. Thus, TSUMUGI can uncover functional gene modules that may remain undetected by knowledge-based interaction networks.

From a future perspective, TSUMUGI will incorporate additional network layers, including PPI and biological pathways, to facilitate mechanistic context for interpreting phenotype-associated gene modules. Furthermore, the inclusion of gene expression profiles will enable the investigation of tissue specificity and inter-organ communication contributing to organism-level traits. Temporal expression dynamics and sex-dependent gene expression will further allow the characterization of life-stage- and sex-specific regulatory programs that drive phenotypic diversity. In addition, leveraging genome-wide association studies and polygenic risk scores may provide complementary insights into the genetic architecture of human diseases. Collectively, these enhancements will expand the scope of TSUMUGI and enhance its utility for dissecting complex traits.

## 5. Conclusion

Here, we present TSUMUGI, a platform for identifying a phenotype-driven gene network using comprehensive KO mouse phenotype data from the IMPC. TSUMUGI enables intuitive exploration of gene relationships associated with organism-level traits through an interactive web interface, alongside flexible and customizable filtering via a command-line interface. By converting gene-phenotype associations into a network representation, TSUMUGI facilitates the discovery of gene interactions underlying organismal phenotypes and human diseases.

## Acknowledgements

We thank Dr. Atsushi Yoshiki, Dr. Shinya Ayabe, Dr. Hiroshi Masuya and Dr. Tatsuya Kushida for their valuable insights and discussions regarding this study. We also thank Dr. Haruka Kuno for her advice on designing the web interface.

## Author contributions

Akihiro Kuno (Conceptualization [lead], Data curation [lead], Formal analysis [lead], Investigation [lead], Methodology [lead], Software [lead], Visualization [lead], Funding acquisition [lead], Writing—original draft [lead], Writing—review & editing [lead]), Kinari Matsumoto (Conceptualization [supporting], Data curation [supporting], Software [supporting], Writing—review & editing [equal]), Taito Taki (Conceptualization [supporting], Software [supporting], Writing—review & editing [equal]), Satoru Takahashi (Conceptualization [supporting], Funding acquisition [supporting], Writing—review & editing [equal]), and Seiya Mizuno (Conceptualization [supporting], Funding acquisition [lead], Writing—review & editing [equal])

