## Supplementary material for "TSUMUGI: a platform for phenotype-driven gene network identification from comprehensive knockout mouse phenotyping data": Fig. S1

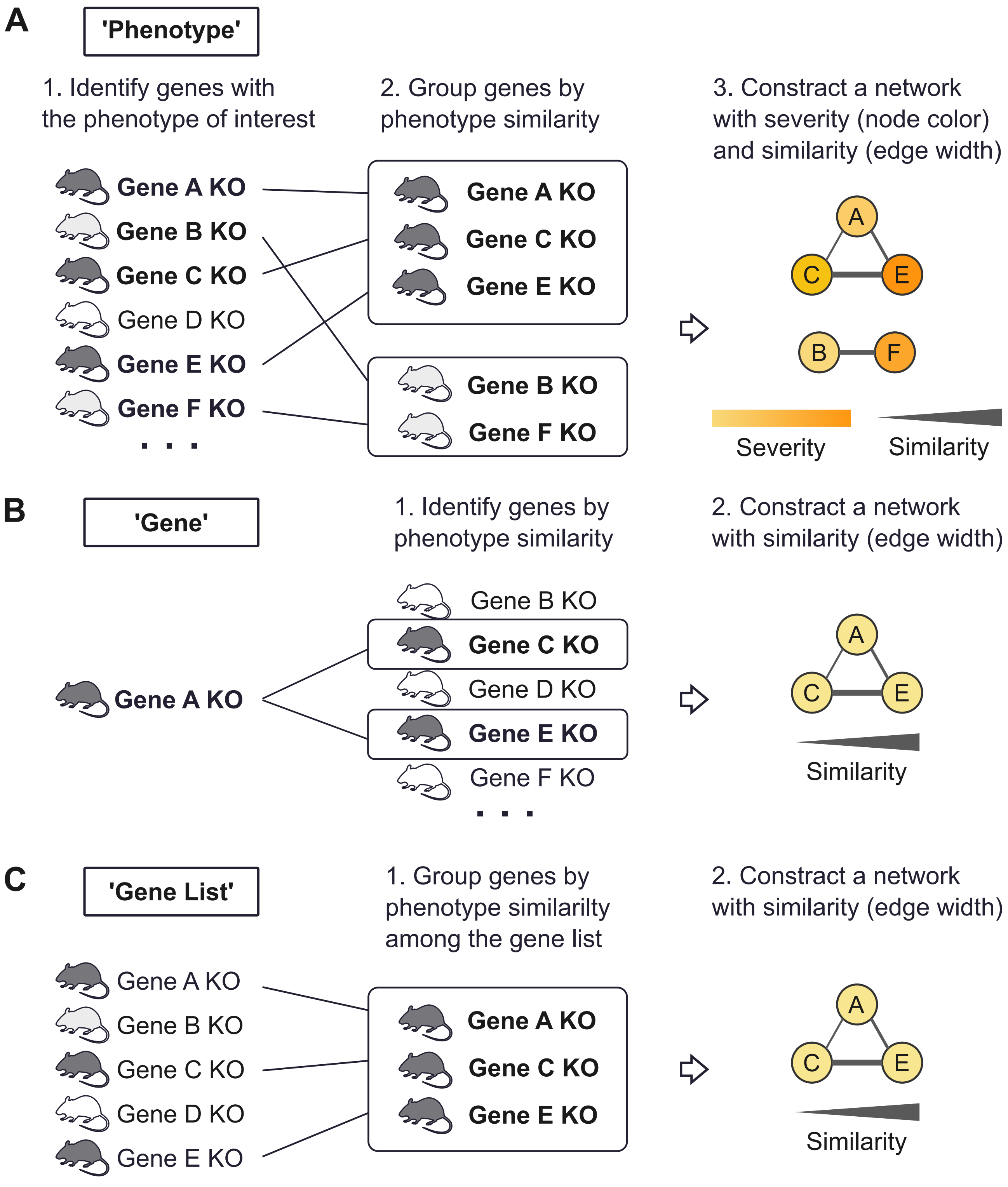


**Supplementary Figure 1.** Three TSUMUGI input types. (A) 'Phenotype' identifies genes whose KO mice exhibit the user-specified phenotype and constructs a phenotype similarity network. Node color reflects phenotype severity when available. (B) 'Gene' identifies genes whose KO mouse phenotypes are similar to those of the user-specified gene. (C) 'Gene List' constructs a phenotype similarity network among the user-specified genes.
