## Supplementary material for "TSUMUGI: a platform for phenotype-driven gene network identification from comprehensive knockout mouse phenotyping data": Fig. S2

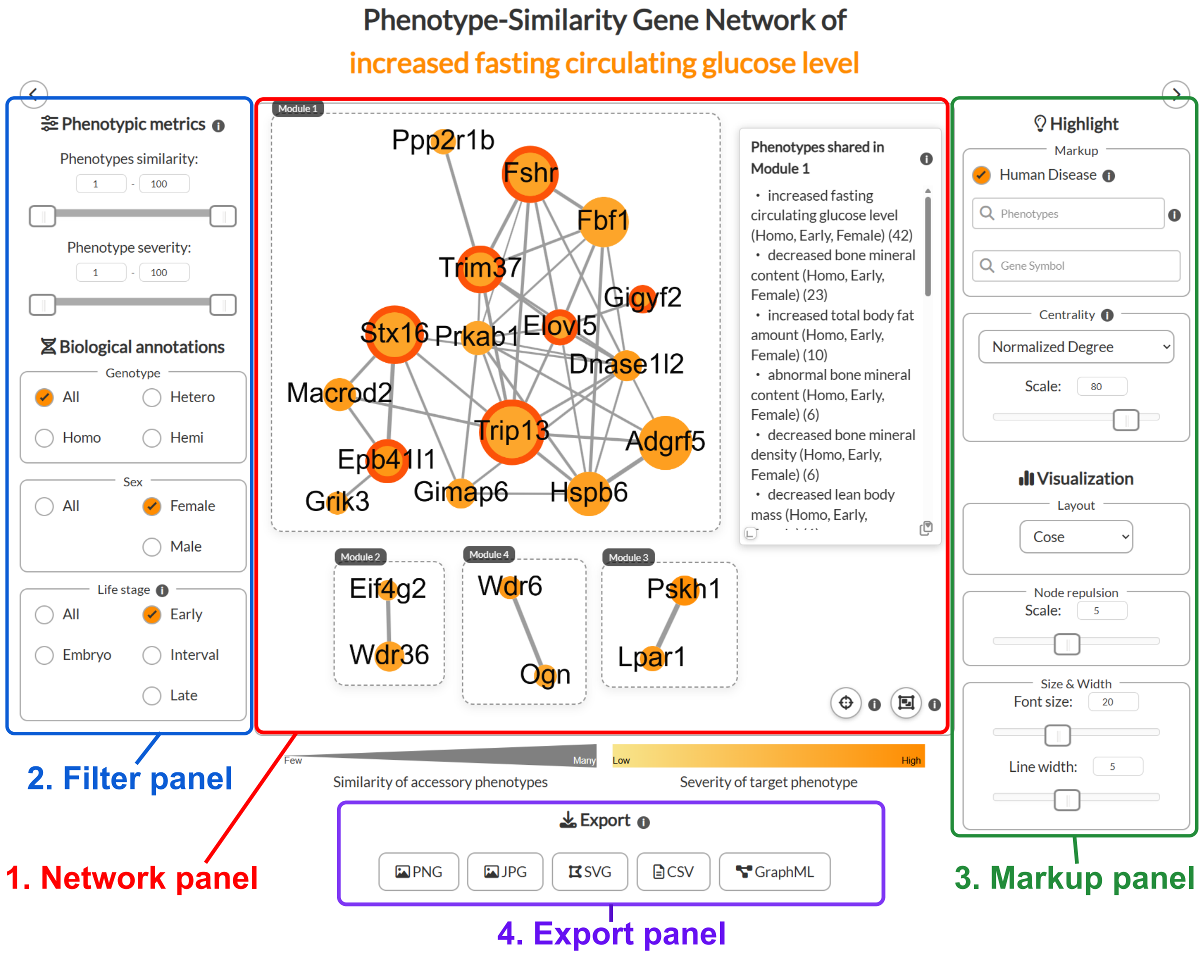


**Supplementary Figure 2.** TSUMUGI network view. The example illustrates a phenotype-similarity gene network for increased fasting circulating glucose levels, restricted to female mice at an early life stage (less than 16 weeks of age).

The network panel (red) displays nodes and edges representing gene symbols and phenotype similarity. Node color indicates phenotype severity, whereas edge width reflects the strength of phenotype similarity. By clicking a module, users can view a list of phenotypes associated with the genes contained within that module.

The filter panel (blue) provides network filtering options based on phenotype similarity, severity, and biological annotations, including genotype, sex, and life stage.

The markup panel (green) offers node-highlighting features, such as red outlines indicating relationships with human diseases, highlighting of phenotypes or genes of interest, and node size adjustment based on centrality measures (degree or betweenness). Additional options include network layout selection, node repulsion control, and text size adjustment.

The export panel enables the network to be exported in multiple formats, including PNG, JPG, SVG, CSV, and GraphML.
